# Polygenic hazard score models for the prediction of Alzheimer’s free survival using the lasso for Cox’s proportional hazards model

**DOI:** 10.1101/2024.04.18.590111

**Authors:** Georg Hahn, Dmitry Prokopenko, Julian Hecker, Sharon M. Lutz, Kristina Mullin, Alzheimer’s Disease Neuroimaging Initiative (ADNI), Rudolph E. Tanzi, Stacia DeSantis, Christoph Lange

**Affiliations:** Harvard T.H. Chan School of Public Health, 677 Huntington Ave, Boston, MA 02115; Genetics and Aging Research Unit, McCance Center for Brain Health, Department of Neurology, Massachusetts General Hospital, Boston, MA 02114; Channing Divsion of Network Medicine, Brigham and Women’s Hospital and Harvard Medical School, Boston, MA 02115, USA; The University of Texas Health Science Center, Houston, TX 77030

**Keywords:** Alzheimer, Cox proportional hazard, Lasso, Penalized regression, Survival

## Abstract

The prediction of the susceptibility of an individual to a certain disease is an important and timely research area. An established technique is to estimate the risk of an individual with the help of an integrated risk model, that is a polygenic risk score with added epidemiological covariates. However, integrated risk models do not capture any time dependence, and may provide a point estimate of the relative risk with respect to a reference population. The aim of this work is twofold. First, we explore and advocate the idea of predicting the time dependent hazard and survival (defined as disease free time) of an individual for the onset of a disease. This provides a practitioner with a much more differentiated view of the absolute survival as a function of time. Second, to compute the time dependent risk of an individual, we use published methodology to fit a Cox’s proportional hazard model to data from a genetic SNP study of time to Alzheimer’s disease (AD) onset, using the lasso to incorporate further epidemiological variables such as sex, APOE (apolipoprotein E, a genetic risk factor for AD) status, ten leading principal components, and selected genomic loci. We apply the lasso for Cox’s proportional hazards to a dataset of 6792 AD patients (composed of 4102 cases and 2690 controls) and 87 covariates. We demonstrate that fitting a lasso model for Cox’s proportional hazards allows one to obtain more accurate survival curves than with state-of-the-art (likelihood-based) methods. Moreover, the methodology allows one to obtain personalized survival curves for a patient, thus giving a much more differentiated view of the expected progression of a disease than the view offered by integrated risk models. The runtime to compute personalized survival curves is under a minute for the entire dataset of AD patients, thus enabling it to handle datasets with 60, 000 to 100, 000 subjects in less than one hour.

## 1 Introduction

Estimating the susceptibility of an individual to a given disease is an important challenge in the age of precision medicine. A widespread approach to provide such an estimate is a polygenic risk score (The International Schizophrenia Consortium, 2009), a measure which quantifies the aggregated genetic risk for a disease or trait. A polygenic risk score with added epidemiological covariates (such as age or sex) is called an integrated risk model (Wand et al., 2021).

Polygenic risk scores and integrated risk models have shown the potential to identify people at high risk for certain diseases in broad-scale clinical settings (Khera et al., 2018; Inouye et al., 2018; Lambert et al., 2019), and they have been developed for a variety of scenarios (Duncan et al., 2019). For instance, they exist for the prediction of the onset of cardiovascular disease (Knowles and Ashley, 2018), type 1 diabetes and coronary artery disease (Mak et al., 2017), or atrial fibrillation (Huang and Darbar, 2017). However, integrated risk models have two major drawbacks. First, they only provide point estimates, and thus do not capture the time dependent risk of an individual. Second, integrated risk models may provide a measure of the relative risk with respect to a reference population (Knowles and Ashley, 2018).

The aim of our contribution is to explore the computation of personalized time dependent hazard and survival curves, computed with an extension of Cox’s proportional hazards under the assumption that censoring and event times are independent. As an application, we focus on Alzheimer’s free survival. Here, the term *Alzheimer’s free survival* is defined as not having contracted Alzheimer’s disease (AD) up to a specific point in time. We are interested in computing personalized hazard (or survival) curves, defined as functions which depend on the genetic and epidemiological characteristics of an individual and return their absolute hazard (or survival) as a function of time. Such hazard and survival curves have the potential to overcome the shortcomings of integrated risk models, in the sense that they provide a practitioner with a much more differentiated view of the absolute risk as a function of time.

Our article is built upon a series of methodological works aimed at applying or extending Cox’s proportional hazards model. Capturing the time dependent risk has been attempted as early as Fraser and Shavlik (1999), who propose a (parametric) multivariate exponential hazard rate model to estimate the lifetime risk and average age at onset of a disease. In Prentice and Kalbfleisch (1979), the authors include a set of fixed covariates into the likelihood for fitting the Cox model through an additional exponential term. This is also summarized in the textbook of Kalbfleisch and Prentice (2002). One year after the seminal publication of Tibshirani (1996), the author also applied the lasso to Cox’s proportional hazards (Tibshirani, 1997), however without the inclusion of additional covariates. The inclusion of covariates into Cox’s proportional hazards model using the adaptive lasso (Zou, 2006) was presented in Zhang and Lu (2007), which we use as the basis of this article. We refer to this method as *Cox-lasso*. Using the Cox-lasso, we fit a Cox’s proportional hazards model to a dataset of 6792 AD patients (composed of 4102 cases and 2690 controls) and 87 covariates; those are age, sex, APOE status, ten leading principal components, and selected genomic loci.

We compare the Cox-lasso to the popular methodology of Desikan et al. (2017) who present a polygenic hazard score for an individual that integrates AD-associated SNPs (single nucleotide polymorphisms) from the International Genomics of Alzheimer’s Project into a Cox’s proportional hazards model together with other covariates. In contrast to the Cox-lasso, the model of Desikan et al. (2017) is based on a conditional partial likelihood and does not incorporate an *L*_1_ penalty to perform model selection. The model of Desikan et al. (2017) has been applied in various works (Tan et al., 2018; Leonenko et al., 2019; Motazedi et al., 2022). Other approaches include the application of the standard Cox model to cardiovascular disease data (Jia et al., 2019), Bayesian quantile regression models (Yang et al., 2019), Cox regression models for competing risks (Gerds et al., 2022; Ozenne et al., 2017), and competing risks in multi-state models (Putter et al., 2021, 2007).

The purpose of our experiments is threefold. First, we demonstrate that the Cox-lasso yields more meaningful predictions of the disease risk as a function of time than classical integrated risk models. Second, we assess the accuracy of the Cox-lasso in comparison with the model of Desikan et al. (2017), showing that the Cox-lasso allows one to obtain survival curves more accurately than with state-of-the-art methods. Third, we demonstrate how to compute personalized survival curves for a patient, thus giving a much more differentiated view on the expected progression of a disease than the view offered by integrated risk models.

The article is structured as follows. The lasso model for Cox’s proportional hazards is discussed in Section 2. Section 3 shows experimental results on the AD dataset. The article concludes with a discussion in Section 4. In the entire article, the indicator function is denoted as 𝕀.

## 2 Methods

This section reviews the Cox-lasso (Zhang and Lu, 2007) which we fit to predict AD free survival (Section 2.1) and discusses some computational considerations of this model (Section 2.2). Moreover, we outline the computation of both survival curves (Section 2.3) as well as survival confidence bands (Section 2.4). The section concludes with a summary of other approaches published in the literature (Section 2.5) which we use in our experimental results.

### 2.1 The Lasso for Cox’s proportional hazards model

The hazard and survival curves we compute are based on the model of Zhang and Lu (2007), who consider a classical Cox’s proportional hazards model (Cox, 1972, 1975) augmented with some covariates *z* ∈ ℝ^*p*^ for fixed *p* ∈ ℕ. It is given by the semiparametric form

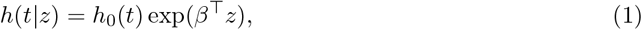

where *h*_0_(*t*) is the time dependent baseline hazard function, and *β* ∈ ℝ^*p*^ is an unknown regression parameter to be fitted. The baseline hazard function has to be estimated on the data.

To fit *β*, Zhang and Lu (2007) propose the following two-step procedure based on a partial log likelihood. Let *n* ∈ ℕ be the number of subjects in the study, and denote with *T*_*i*_ and *C*_*i*_ the failure time and censoring time of individual *i* ∈ {1, …, *n*}, respectively. For each individual *i* ∈ {1, …, *n*}, we assume that we have additional covariates *z*_*i*_ ∈ ℝ^*p*^. First, defining δ_*i*_ = 𝕀(*T*_*i*_ ≤ *C*_*i*_), the following partial log likelihood is fitted to the data under investigation:

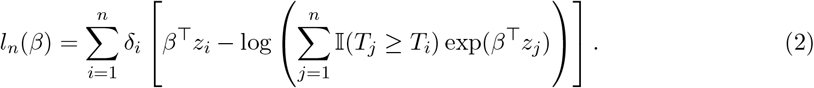

In eq. (2), each individual whose failure time *T*_*i*_ occurred before the censoring time *C*_*i*_ contributes to the log likelihood. The log likelihood itself consists of a regression term *β*^⊤^*z*_*i*_ for each individual *i* ∈ {1, …, *n*}, adjusted (that is, normalized) by the contributions of the other individuals *j* with failure times *T*_*j*_ ≥ *T*_*i*_. Note that the normalization becomes a subtraction after taking logarithms. Second, using eq. (2), Zhang and Lu (2007) propose an adaptive lasso estimator (Zou, 2006) for *β* given by

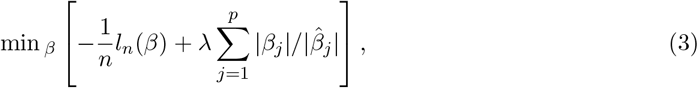

where the entries 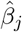 for *j* ∈ {1, …, *p*} stem from the maximizer 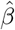 of the partial log likelihood of eq. (2) which was computed first. Due to the fact that 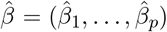 are consistent estimators, their assigned values reflect the importance of the covariates, which in turn serves as a weighting scheme in eq. (3), see Zou (2006). The lasso parameter *λ >* 0 in eq. (3) controls the level of sparsity as usual, and it is determined in this work using cross validation (Hastie et al., 2016).

### 2.2 Computational considerations

To minimize eq. (2) and thus fit the values 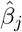 for *j* ∈ {1, …, *d*}, we employ the *optim* function in R using the *BFGS* algorithm (Broyden, 1970; Fletcher, 1970; Goldfarb, 1970; Shanno, 1970) with numerical gradients. Alternatively, the simulated annealing algorithm (Kirkpatrick et al., 1983) can be used, available in the *optim* function under the name *SANN*. For numerical stability, we norm the resulting coefficients of the vector 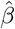 to have an *L*_2_ norm of one.

After obtaining 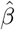 from eq. (2), eq. (3) is likewise minimized for *β* with the help of the *optim* function in R. The fitted coefficients 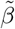 then stay invariant for the population that was being fitted. For any individual *i*, for instance in a validation set *i* ∈ *V* ⊂ {1, …, *n*}, we obtain a fully specified Cox-lasso model via eq. (1) as *h*_*i*_(*t*) = *h*_0_(*t*) exp 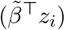 where *h*_0_(*t*) is the baseline hazard fitted separately, 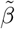 is the minimizer of eq. (3), and *z*_*i*_ is the vector of covariates for individual *i*.

We compute an estimate of the baseline hazard *h*_0_(*t*) with the help of the function *basehaz* of the *survival* package on CRAN (Therneau et al., 2022). The baseline hazard is defined as

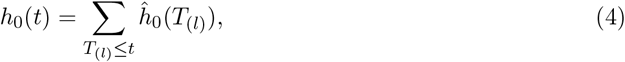

with the functions 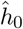 being defined as

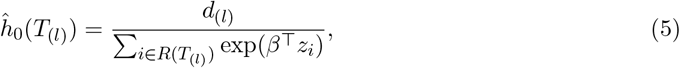

where *T*_(1)_ < *T*_(2)_ < ⃛ < *T*_(*n*)_ denote the ordered event times, *d*_(*l*)_ is the number of events at time *T*_(*l*)_, and *R*(*T*_(*l*)_) is the set containing all individuals still susceptible to the event at *T*_(*l*)_. The function *basehaz* returns a numerical representation of the baseline hazard *h*_0_(*t*) at a grid of time points.

As it is common practice, we will employ a classical training and validation setup in our experimental results of Section 3.

### 2.3 Computation of the survival curve

The expression of eq. (1) gives a full specification of the time dependent hazard function *h*(*t*) := *h*(*t*|*z*). Using the relationship 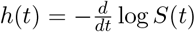 of the hazard function and the survival function *S*(*t*), it can be seen that the survival function can be expressed as

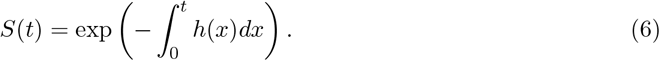

Therefore, to calculate the survival function *S*(*t*) for a given time *t* from the hazard function, we numerically integrate the hazard function from 0 to *t*.

### 2.4 Computation of confidence bands for the hazard and survival functions

We are interested in quantifying the uncertainty in the computation of the hazard function *h*(*t*) and the survival function *S*(*t*) for a given individual. This can be done as follows using a bootstrapping approach. For a given training and validation split, let *n*_0_ < *n* denote the number of individuals in the training set. Let *B* ∈ ℕ be the number of bootstrap samples chosen by the user. We repeatedly draw *B* training sets *σ*_1_, …, *σ*_*B*_ ⊂ {1, …, *n*} with replacement. Using the individuals in each training set *σ*_*i*_ only, where *i* ∈ {1, …, *B*}, we compute the hazard function *h*(*t*) and survival function *S*(*t*) as outlined in Section 2.2 and Section 2.3. For each time point *t*, this strategy thus results in *B* estimates of the hazard and survival, respectively, which immediately allows one to compute an empirical lower (for instance, 5%) and upper (for instance, 95%) confidence bound for *t*.

### 2.5 Other approaches

In the experiments of Section 3, we compare the Cox-lasso to other state-of-the-art methods. This section gives a brief overview of them.

In Desikan et al. (2017), the authors aim to predict the AD age of onset using a forward stepwise Cox regression with conditional partial likelihood. Using our notation of Section 2.1, they define

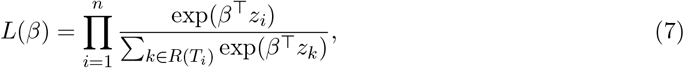

where *R*(*T*_*i*_) is the set of all individuals *j* ∈ {1, …, *n*} whose time point *T*_*j*_ (in our case, the age) is less or equal than *T*_*i*_. The vectors *z*_*i*_ per individual *i* ∈ {1, …, *n*} encode all covariates. We optimize *L*(*β*) for *β* using the *optim* function in R, and normalize the resulting coefficients to have an *L*_2_ norm of one as done in Section 2.2. To obtain a time dependent hazard curve *h*(*t*), we estimate the baseline hazard as done in Section 2.2 and multiply it with an exponential term as done in eq. (1) to obtain a fully specified hazard function. The survival curve was obtained from the hazard curve via numerical integration as described in Section 2.3.

Cox regression models for competing risks (Gerds et al., 2022; Ozenne et al., 2017), and competing risks in multi-state models (Putter et al., 2021, 2007) are presently unconsidered in this work. The function *Survfit*, provided in the *survival* R-package on CRAN (Therneau et al., 2022), is unconsidered as it is not supposed to run genomic level data.

When comparing the Cox-lasso to classical integrated risk models, we employ the *lassosum* R-package (Mak et al., 2020) which implements the methodology of Mak et al. (2017).

## 3 Results

In this section, we present numerical results on the Cox-lasso of Section 2 applied to a dataset of AD onset. We start by introducing the dataset under consideration (Section 3.1). In Section 3.2 we examine the hazard and survival curves obtained with the Cox-lasso and the model of Desikan et al. (2017). We compare our predicted survival curves with the ones obtained by using a regular integrated risk model in Section 3.3. Section 3.4 evaluates the Cox-lasso with respect to its accuracy. We conclude with the computation of confidence bands for personalized (that is, per patient) AD hazard and AD survival forecasts in Section 3.5.

### 3.1 The Alzheimer’s dataset

We have used a whole genome sequencing dataset and harmonized AD affection status from cohorts sequenced by the Alzheimer’s Disease Sequencing Project (ADSP) and other AD and Related Dementia’s studies (Beecham et al., 2017). Vcf files were obtained from the National Institute on Aging Genetics of Alzheimer’s Disease Data Storage Site (NIAGADS) under the accession number *NG00067*.*v5*. We have excluded subjects based on genotype missingness, observed versus expected homozygous genotype counts, and kinship coefficient (first and second degrees of relatedness). Next, we have extracted a subset of non-Hispanic white subjects (NHW) based on self-reported population. This was compared with assignment based on genetic principal components (PC) using the Jaccard matrix on rare variants (Prokopenko et al., 2016) and PC outliers (based on 10 PCs) were removed (i.e., 5 standard deviations away from the mean). From available whole genome sequencing data we have selected 74 SNPs associated with AD (Bellenguez et al., 2022, Supplementary Table 32).

The final dataset consists of 6792 AD patients (composed of 4102 cases and 2690 controls) with full information on 87 covariates. Those covariates are age, sex, APOE status, the first 10 principal components, and the 74 SNPs associated with AD of Bellenguez et al. (2022). The APOE status of each individual was encoded with 1 (single e4), or 2 (e4/e4). The age is recorded as the patient’s age at onset (for cases) or age at last examination (for controls). Both cases and controls are considered together when fitting each model.

### 3.2 Hazard and survival curves

We start by examining the hazard and survival curves obtained by fitting the approach of Section 2.1. We divide the dataset of Section 3.1 into three splits, tuning, training, and validation. To tune the methods, we employ one third of the data or 2264 randomly selected individuals. Using 10-fold cross-validation we tune the lasso penalty of eq. (3) to be *λ* = 0.4. Details on the 10-fold cross-validation can be found in Section A.

The remaining two thirds of the data (4528 individuals) are used for the following results. First, we use a training set of 4000 randomly selected individuals (of the 4528 individuals) to fit the Cox-lasso with penalty *λ* = 0.4 (see Section 2.2) or the model of Desikan et al. (2017) (see Section 2.5). We then plot the hazard and survival curves of the remaining 528 individuals in the validation set. This is for visualization purposes only, since plotting too many hazard and survival curves in the validation set proved to render the figures illegible.

Figure 1 shows the hazard (left) and survival (right) curves for the individuals in the validation set. We observe that the curves seem to reasonably model the AD hazard and survival, in the sense that the hazard increases with age, and the survival (defined as not developing AD) decreases to zero as a function of age. Moreover, we observe that the curves for different individuals display a wide variety of functional shapes. This is a property one would hope for, as it demonstrates that in our model, the genomic and epidemiological data for each individual indeed determines their survival.

**Figure 1.**
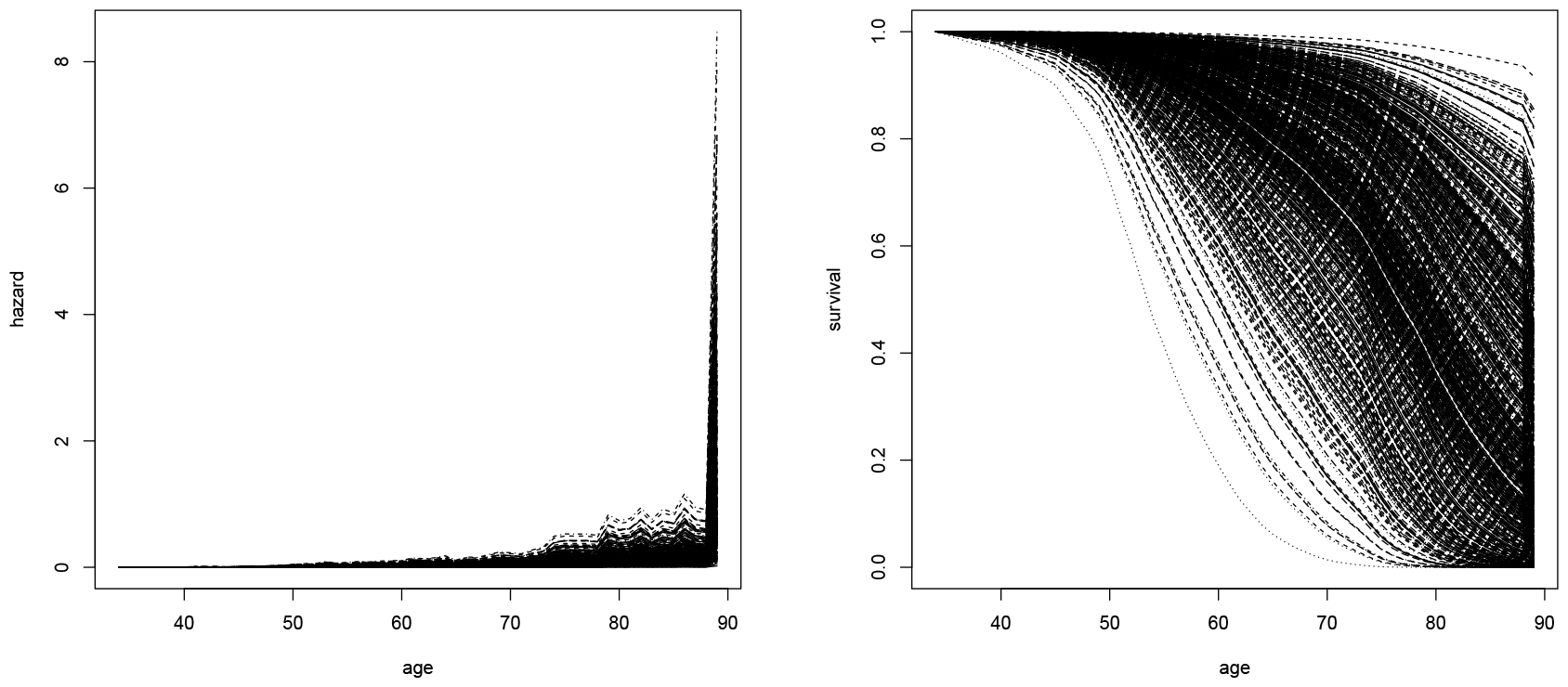
Hazard curves (left) and corresponding survival curves (right) for the Cox-lasso, see Section 2.1. Training set used to calibrate the lasso model, the curves for the individuals in the validation set are displayed.

Figure 2 displays the corresponding hazard and survival curves computed with the help of the model of Desikan et al. (2017) on the same training and validation datasets. We observe that for the Desikan et al. (2017) model, the computed hazards are up to an order of magnitude higher, though the overall functional shape of the hazard and survival curves matches the one of Figure 1.

**Figure 2.**
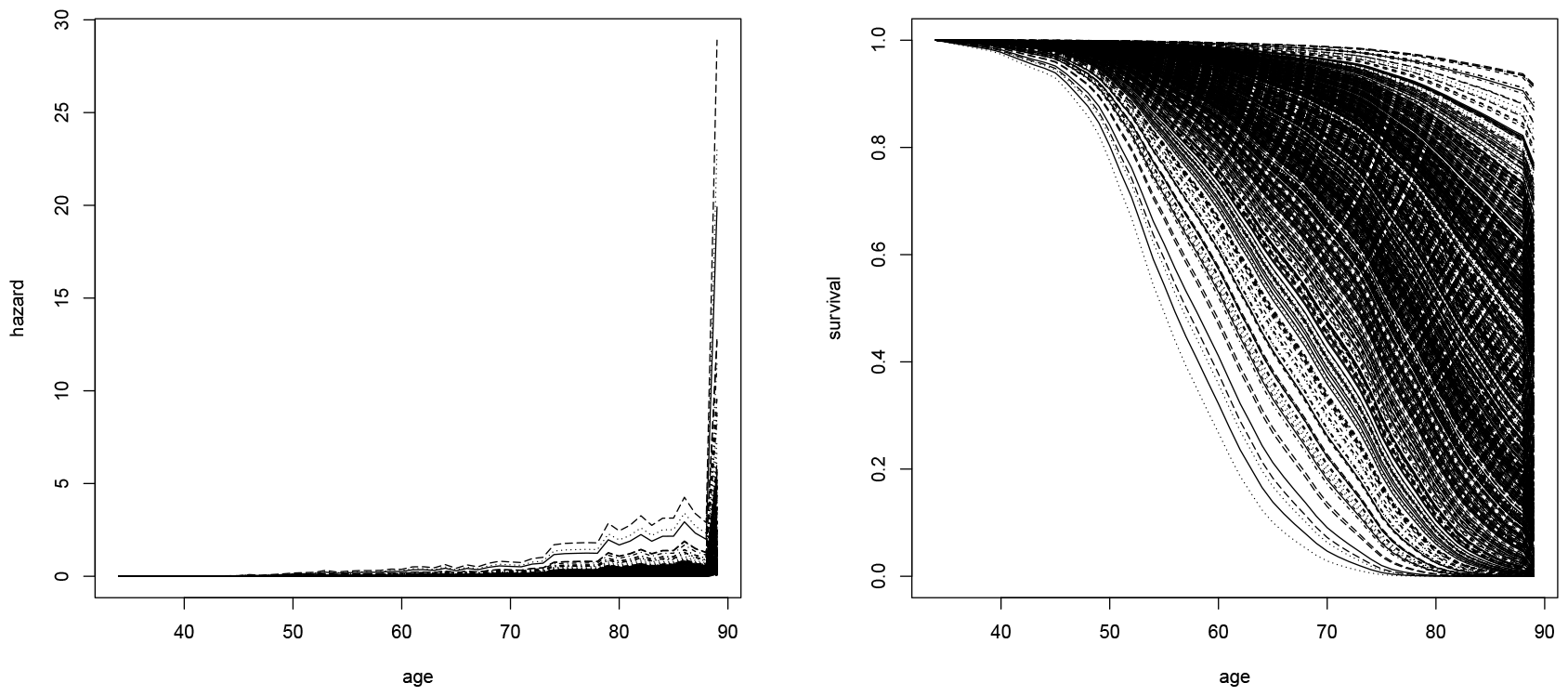
Hazard curves (left) and corresponding survival curves (right) for Desikan et al. (2017). Training and validation sets as in Figure 1.

### 3.3 Comparison with integrated risk models computed as a function of age

We contrast the survival curves presented in Section 3.2 with curves obtained using a classical integrated risk model. In particular, we use *lassosum* of Mak et al. (2017) to fit an integrated risk model to the same training set of Section 3.2, where the AD status is the outcome, and the other 87 covariates are used as predictors.

After having fitted a score on the training set, we use the score to predict the risk for the individuals in the validation set. However, we do not use the actual age of the patients in the validation set. Instead, we vary their ages simultaneously from 50 to 100, thus allowing us to obtain an integrated risk model as a function of age for each individual. In order to interpret each score obtained in this fashion, we compare it with the distribution of scores in the training set. If the score for an individual for any given age is above the mean of the distribution of the training set, we classify the individual as high risk (survival prediction of 0), and otherwise as low risk (survival prediction of 1). By varying the age, we obtain a risk curve for each individual.

Figure 3 shows the curves obtained in this fashion on the validation set. We observe that all individuals seem to transition from the low to the high risk regime at a more narrow age range (between approximately 75 and 85 years of age) than the ones in Figure 1 and Figure 2. This seems unlikely, and it gives much less detailed information for practitioners than the hazard and survival curves displayed in Section 3.2.

**Figure 3.**
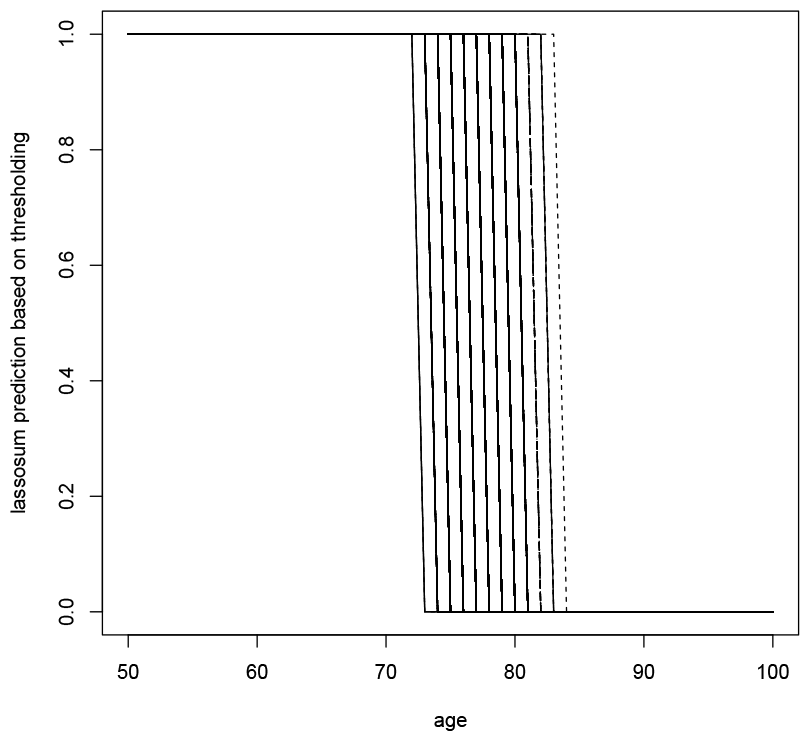
Prediction of lassosum (after thresholding) as a function of age.

Overall, we observe that the Cox-lasso allows one to compute individualized survival curves for the onset of AD which carry more detailed information about the projected course of a disease than the mere point estimates returned by integrated risk models.

### 3.4 Evaluation of two approaches

We aim to evaluate the predictions made by the Cox-lasso and the model of Desikan et al. (2017). To this end, we utilize a modified experimental setup consisting of a training and validation split of the 4528 withheld individuals of Section 3.2 which were not used to tune the lasso. To be precise, we split these individuals into randomly drawn training sets having a size ranging from 100 to 4000 in steps of 100. In each step, the remaining individuals form the validation set. Both the Cox-lasso and the model of Desikan et al. (2017) are trained on the same training set and evaluated on the same validation set. All results presented in this section are averages of 100 repetitions.

For all individuals in the validation set, we know both their AD status and age, where the age is either the most recent follow-up time point for controls, or the age of onset for AD cases. Therefore, for all individuals in the validation set, we first compute their predicted survival curves and then query the survival probability at the age recorded in the validation set. For the controls, a higher prediction of their survival probability is desired (at the age of the last follow-up point). For the cases, a lower survival prediction is better (since we already know that the age describes the age of onset). We consider two metrics. First, we compute the absolute (that is, *L*_1_) loss between the predicted survival probabilities at the recorded ages with the true binary AD outcomes (0 for controls and 1 for cases) and normalize by the number of individuals in the validation set. This measure is always within the interval [0, 1]. Second, we compute the AUC (area under the curve) measure for the continuous survival prediction versus the binary true AD outcomes.

Figure 4 shows the results of this experiment, where a smaller *L*_1_ loss and a higher AUC indicate a better performance. As expected, we observe that for both the Cox-lasso and the Desikan et al. (2017) model, the prediction accuracy increases with the size of the training set used to fit the models. The Cox-lasso seems to exhibit a slightly better performance on our dataset with respect to both the *L*_1_ norm and the AUC. For both the Cox-lasso and the Desikan et al. (2017) model, fitting the hazard score model to the full training set takes around 50 seconds on a standard Intel Core i5-7200U with four CPUs, each having a clock speed of 2.50 GHz and 7.6 GiB of RAM memory. This allows one to handle datasets with 60, 000 to 100, 000 subjects in less than one hour.

**Figure 4.**
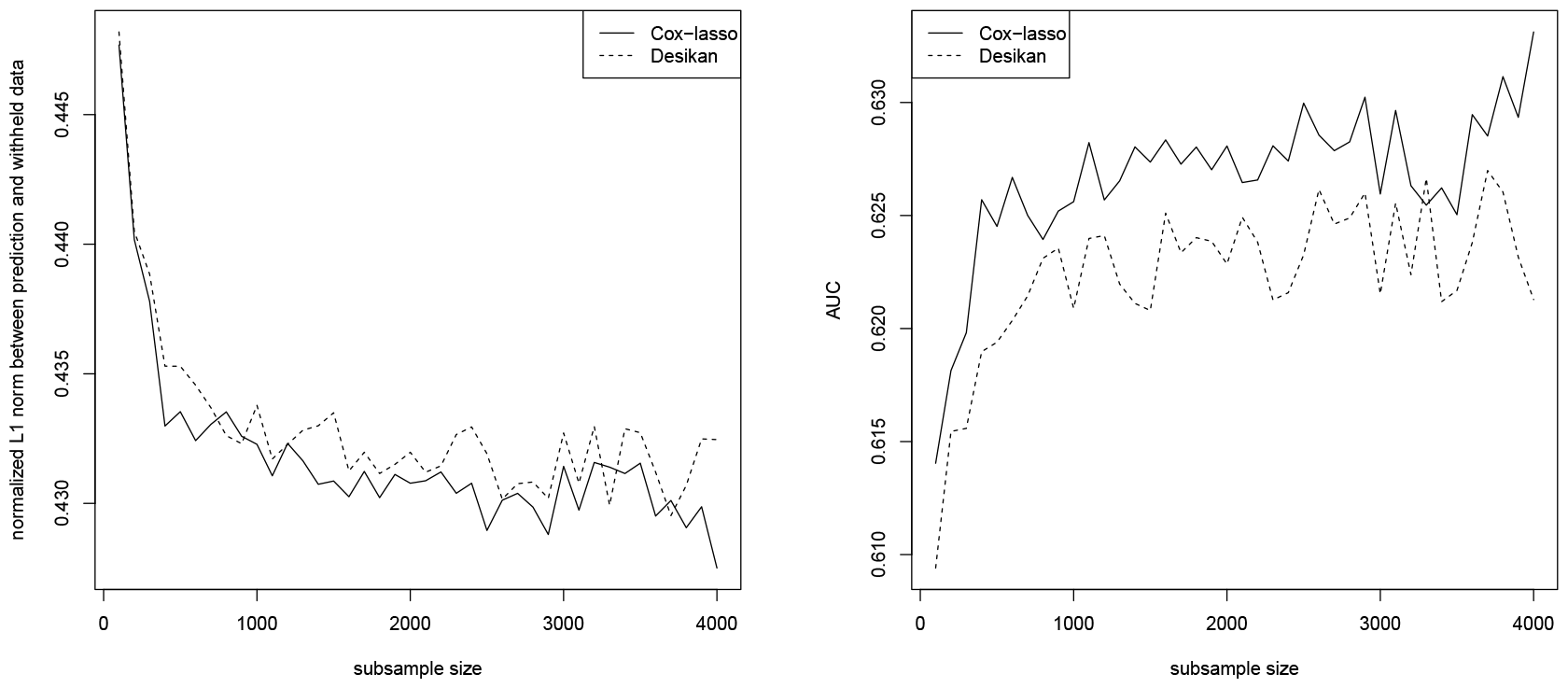
Normalized sum of *L*_1_ loss between predicted survival probability and true AD outcome (left) as well as AUC (right) as a function of the size of the training set. Cox-lasso of Section 2.1 (solid) and model of Desikan et al. (2017) (dashed).

### 3.5 Computation of confidence intervals for individual AD hazard and survival

Finally, we also consider the computation of confidence bands around the predicted hazard and survival curves of the Cox-lasso. These are easily computed using a bootstrap technique as described in Section 2.4. We use 100 bootstrap samples, each time computed on a training set of 1000 randomly selected individuals.

Figure 5 displays the result of the bootstrap procedure for two patients in the dataset of Section 3.1. We observe that indeed, the bootstrap technique allows one to efficiently compute a measure of confidence around the hazard and survival curves. As expected, the hazard confidence bands are narrower at younger ages, and show higher degrees of uncertainty at advanced ages. The two patients displayed in the top and bottom row of Figure 5 exemplarily demonstrate that the shape of the hazard and survival curves is roughly the same for all patients, though the magnitude of the effects (displayed on the y-axis) varies. For instance, the first patient has a much higher AD-free survival (top right) than the second one (bottom right).

**Figure 5.**
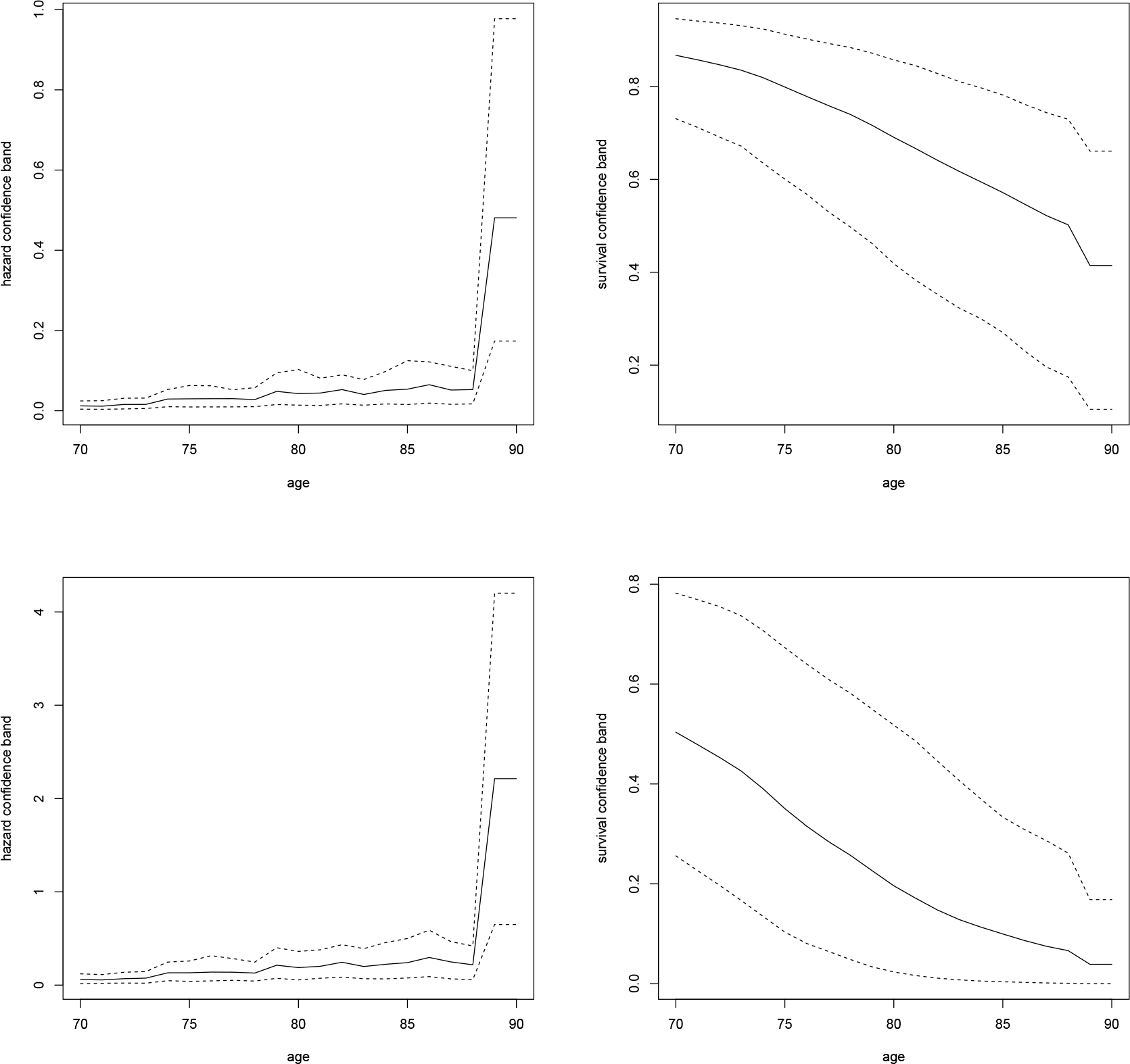
Bootstrap confidence bands for the hazard (left) and survival (right) curves of the Cox-lasso. Patient at lower risk (top row) and higher risk (bottom row). Mean hazard and survival displayed as solid line, and 5% and 95% confidence bands displayed as (lower and upper) dashed lines.

## 4 Discussion

In the age of precision medicine, the prediction of the susceptibility of an individual to certain diseases is a timely and important area of research. The established methodology to compute such an aggregated risk estimate for an individual based on its epidemiologic and genetic traits is an integrated risk model (a polygenic risk score with added epidemiological covariates). However, such integrated risk models do not capture the time dependent risk of an individual. Moreover, they may only provide a measure of the relative risk with respect to a reference population.

In this publication, we focus on a more involved task, the computation of hazard and survival curves. A survival curve indicates the “survival” from the onset of a disease as a function of time (age). We investigate the potential of the lasso for Cox’s proportional hazards (herein called Cox-lasso), introduced in Zhang and Lu (2007), to compute survival predictions for any given individual, using its epidemiological and genetic characteristics. Using an application of Alzheimer’s disease, we demonstrate that the Cox-lasso indeed allows a practitioner to obtain a refined prediction of the progression of AD. Moreover, the Cox-lasso achieves a higher accuracy (measured with the AUC metric) than the popular model of Desikan et al. (2017), and yields more interpretable survival curves than the stepwise curves returned by polygenic risk scores. Due to its simplicity, the runtime of the proposed Cox-lasso is under a minute for the entire dataset of AD patients, thus enabling it to handle datasets with 60, 000 to 100, 000 subjects in less than one hour.

## Acknowledgments

Data collection and sharing for this project was funded by the Alzheimer’s Disease Neuroimaging Initiative (ADNI) (National Institutes of Health Grant U01 AG024904) and DOD ADNI (Department of Defense award number W81XWH-12-2-0012). ADNI is funded by the National Institute on Aging, the National Institute of Biomedical Imaging and Bioengineering, and through generous contributions from the following: AbbVie, Alzheimer’s Association; Alzheimer’s Drug Discovery Foundation; Araclon Biotech; BioClinica, Inc.; Biogen; Bristol-Myers Squibb Company; CereSpir, Inc.; Cogstate; Eisai Inc.; Elan Pharmaceuticals, Inc.; Eli Lilly and Company; EuroImmun; F. Hoffmann-La Roche Ltd and its affiliated company Genentech, Inc.; Fujirebio; GE Healthcare; IXICO Ltd.; Janssen Alzheimer Immunotherapy Research & Development, LLC.; Johnson & Johnson Pharmaceutical Research & Development LLC.; Lumosity; Lundbeck; Merck & Co., Inc.; Meso Scale Diagnostics, LLC.; NeuroRx Research; Neurotrack Technologies; Novartis Pharmaceuticals Corporation; Pfizer Inc.; Piramal Imaging; Servier; Takeda Pharmaceutical Company; and Transition Therapeutics. The Canadian Institutes of Health Research is providing funds to support ADNI clinical sites in Canada. Private sector contributions are facilitated by the Foundation for the National Institutes of Health (www.fnih.org). The grantee organization is the Northern California Institute for Research and Education, and the study is coordinated by the Alzheimer’s Therapeutic Research Institute at the University of Southern California. ADNI data are disseminated by the Laboratory for Neuro Imaging at the University of Southern California.

The computations in this paper were run in part on the FASRC Cannon cluster supported by the FAS Division of Science Research Computing Group at Harvard University. Please refer to the Supplementary Note for full acknowledgements.

## Data availability

The ADSP WGS data set is available from DSS NIAGADS under accession number NG00067.v5.

## A Cross-validation to determine the lasso penalty

We use the setting of Section 3.2 to calibrate the Cox-lasso via 10-fold cross-validation. This is done as follows. Assume the lasso parameter *λ* is fixed. We divide the subset of the data that was allocated to tuning (consisting of 2264 randomly selected individuals, see Section 3.2) into k = 10 (randomly drawn) folds. We apply the Cox-lasso for the given *λ* to this data with each fold excluded once. Each time, we evaluate the obtained fit on the withheld fold, using the measures outlined in Section 3.4. The measure of accuracy we employ is the *L*_1_ norm between predicted survival probabilities and the true AD outcome of the withheld fold. This yields k = 10 measurements of the *L*_1_ norm (one for each excluded fold) for each fixed choice of *λ*.

The lasso parameter *λ* of eq. (3) is now varied in the interval [0, 1] in steps of 0.2. For each value of *λ*, we average the *k* = 10 measurements of the *L*_1_ measure (one for each fold). The averages are plotted as a function of *λ* in Figure 6. Since we aim to minimize the *L*_1_ prediction error on the withheld folds, based on Figure 6, we choose *λ* = 0.4 for the Cox-lasso in our experiments of Section 3.

**Figure 6.**
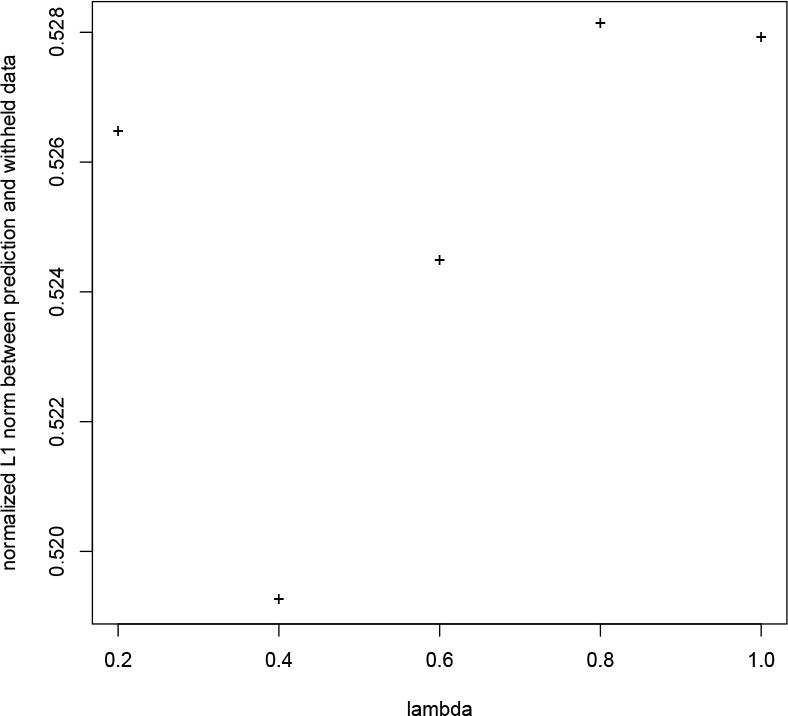
Cross-validation results showing the normalized *L*_1_ error as a function of *λ* ∈ [0, 1].

